# Evidence of optimal control theory over active inference in corticospinal excitability modulations

**DOI:** 10.1101/2024.05.06.592675

**Authors:** I.M. Brandt, T. Grünbaum, M.S. Christensen

**Affiliations:** Department of Neuroscience, University of Copenhagen, Denmark; Department of Psychology, University of Copenhagen, Denmark; Section for Philosophy, University of Copenhagen, Denmark; CoInAct Research Group, University of Copenhagen, Denmark

**Keywords:** Active inference, optimal control theory, motor command, proprioceptive prediction error, motor evoked potential (MEP)

## Abstract

Two theories, optimal control theory and active inference, dominate the motor control field. We use transcranial magnetic stimulation (TMS) in force and angle tasks to examine whether corticospinal excitability represents a motor command, as proposed by the optimal control theory, or a proprioceptive prediction, as proposed by active inference. Our results strongly support optimal control theory. We encourage comparisons of the theories against each other based on empirically testable predictions.

## Main

Two prominent theories are competing to explain motor control: *optimal control theory* and *active inference*. Both theories are theoretically convincing, but it has proved hard to distinguish them empirically. Experimentally testable hypotheses comparing the two theories have to our knowledge not been formulated. Here we present a novel experimental setup with the aim to test these two leading motor control theories against each other. We encourage proponents of each theory to formulate experimentally testable hypotheses to allow the field to move forward based on empirical studies.

According to optimal control theory, to create movement, the brain uses an inverse model to convert a desired movement goal into movement signals called motor commands (Fig. 1). Motor commands are motor signals that descend from the primary motor cortex (M1) propagating via the corticospinal tract, activating alpha motor neurons (αMN), ultimately innervating target muscles. Shadmehr et al (2010) present the motor command as a motor signal conveying information that reflects the current state of the movement^1^. ‘The current state’ can be interpreted in multiple ways. In this paper we use it in terms of current parameter measures of either force (normalized to maximal voluntary contraction (MVC)) or joint angle (e.g. position).

**Figure 1.**
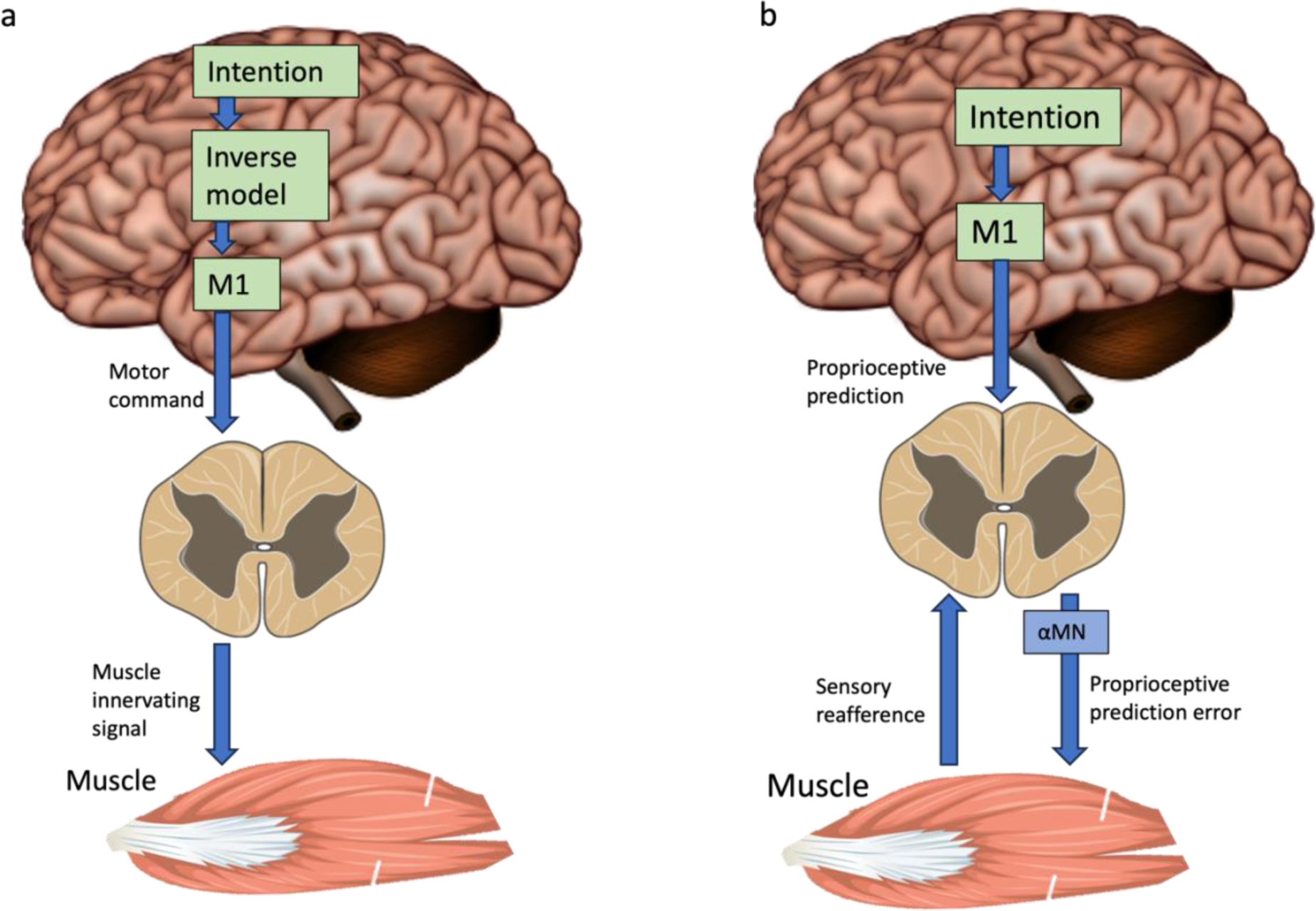
Optimal control theory and active inference. **a) shows relevant signals for** optimal control theory, **b)** shows relevant signals for active inference. For optimal control theory, an intention is through the inverse model changed into a motor plan that is executed through motor commands propagating from M1 through the corticospinal tract to the target muscle. In optimal control theory, it is thus a motor command that activates the muscle. For active inference, the intention is converted into proprioceptive predictions. These predictions originate from M1 and meet sensory reafference in either the spinal dorsal horn, dorsal column nuclei in the brain stem, or in the thalamus. The difference between proprioceptive predictions and sensory afference is the proprioceptive prediction error, which propagate through the αMN to the target muscle. According to active inference, it is thus the prediction error that activates the muscle. Figure modified from^23–25^.

Active inference builds upon a statistical physics principle of minimizing free energy^2,3^, which means minimizing surprise by reducing prediction error. According to active inference, the descending signals from M1 are proprioceptive predictions rather than motor commands, i.e. predictions about which proprioceptive signals the movement should induce (Fig. 1). Thus, according to active inference, the outputs from M1 are sensory in nature, contrasting with the motor-centric view of optimal control theory. The proprioceptive predictions descending from M1 are transformed into proprioceptive prediction errors upon encountering ascending sensory information either in the spinal cord, in the brain stem, or in the thalamus^4^. Therefore, the αMN carry a signal that corresponds to the difference between the predicted proprioceptive feedback and the actual ascending sensory signal, i.e., the prediction error. According to active inference, as presented by Adams et al (2013), it is this prediction error that drives the movement^4^.

To test the theories against each other, we developed a study with force and angle tasks. Thirty-three participants completed the study and signed written informed consents. The protocol was approved by the ethics committee of The Capital Region of Denmark (H-21061035) and data collection and storage was performed following UCPH guidelines (2004334 – 4242). All participants performed both tasks (Fig. 2). We used transcranial magnetic stimulation (TMS) induced motor evoked potentials (MEPs) during movements to measure the change in corticospinal excitability (CSE)^5,6^. MEP amplitude is modulated by state changes in CSE and can be used as a read-out of the signal propagating from M1 through the corticospinal pathway to the motoneurons, ultimately innervating target muscles. In both tasks, participants were sitting in front of a table with the forearm placed in a semiprone position, palm facing medially and without visual feedback of their movement. In the force task, participants performed isometric index finger force production at different force levels. They were stimulated with TMS at different force levels toward the maximal force level in a trial. The angle task was a planar flexion-extension index finger movement task, moving the metacarpophalangeal joint. Participants performed different angle levels and were stimulated at different angles (Fig. 2, see online Methods).

**Figure 2.**
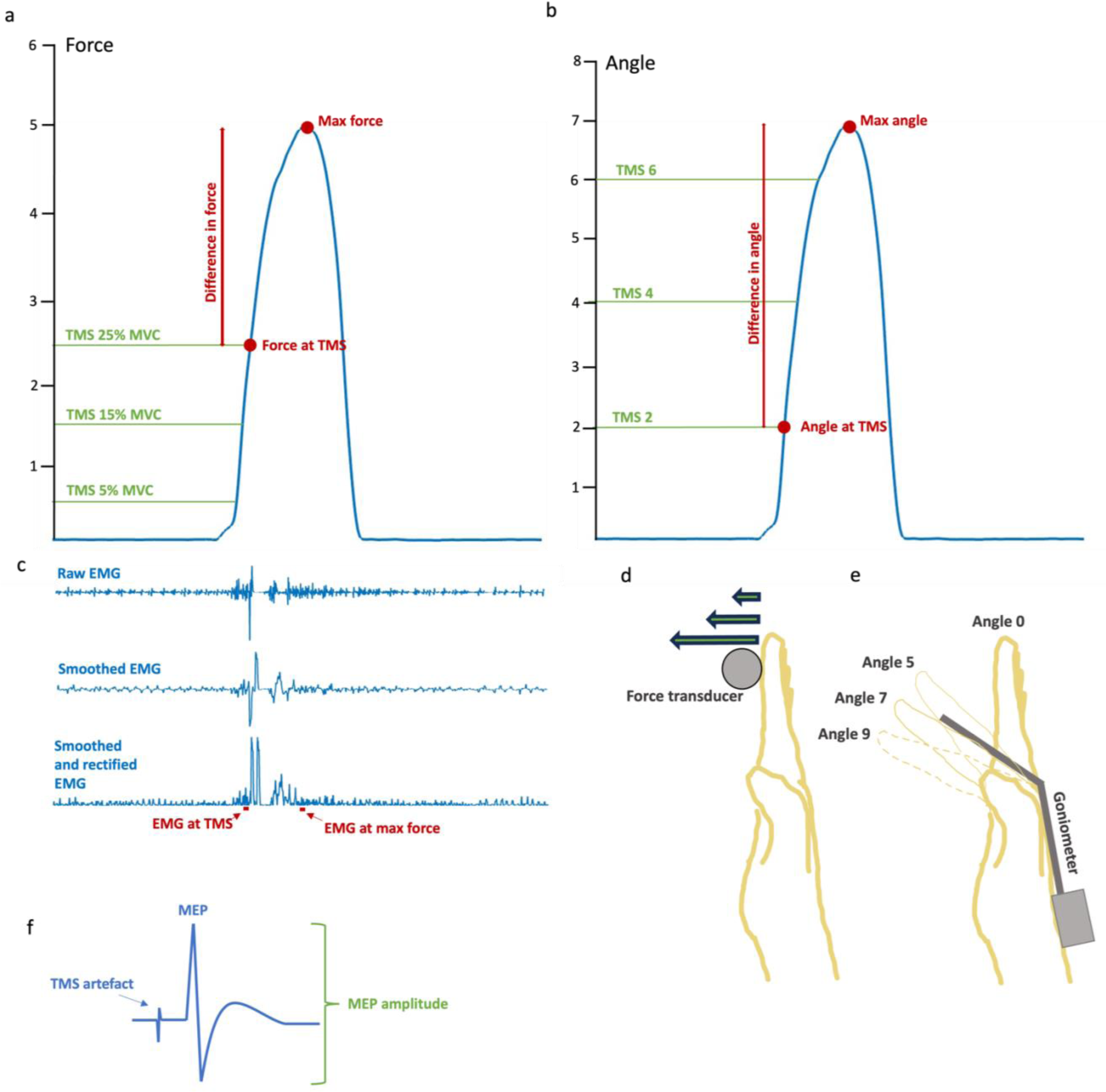
Experimental setup. A) Force task. The blue line signifies the force measured with the force transducer. The green lines indicate when a TMS stimulation could occur, only one stimulation occurred in each trial. In the force task, the participant aimed for either force level 1, 3, or 5. When aiming for force level 1 (10% MVC), the participant was stimulated at 5% MVC. When aiming for force level 3 (30% MVC), the participant could be stimulated at 5% or 15% MVC. When aiming for force level 5 (50% MVC), the participant could be stimulated at 5%, 15% or 25% MVC. Force at TMS and difference between force at TMS and max force was measured. B) Angle task. The blue line signifies the goniometer measure of angle. The green lines indicate when a TMS stimulation could occur, only one stimulation occurred in each trial. In the angle task, the participant aimed for either angle 5 or 7. When aiming for angle 5, the participant could be stimulated at either angle 2 or 4, when aiming for angle 7, the participant could be stimulated at either angle 2, 4, or 6. In this example, participant performed angle 7 and was stimulated at angle 2. Angle at TMS and difference between angle at TMS and max angle was measured. C) Example of raw EMG, smoothed EMG, and smoothed and rectified EMG. EMG at TMS is the mean smoothed, rectified EMG 55-5ms prior to TMS stimulation. EMG at max force is the mean smoothed, rectified EMG 50-0ms prior to max force. D) Hand position in the force task. Right index finger pressed isometrically on force transducer. E) Hand position in the angle task. A goniometer measuring the angle was attached to the right index finger. F) Example of MEP and MEP amplitude. See online Methods section for further details.

According to optimal control theory, without perturbations to the movement, the magnitude of the motor command should be proportional to the state of the produced movement^1^. That is, absent perturbation, the motor command and thus the MEP should reflect the current state. Therefore, the model representing optimal motor control theory includes the variable force at TMS or angle at TMS.

According to active inference, the descending signal is a proprioceptive prediction that is turned into a prediction error. This entails that both prediction and prediction error should be reflected in the MEP amplitude. Since the prediction should remain constant, our analysis prioritizes the prediction error, which reflects the difference between the current state and the end-state of a movement. In the model representing active inference, we therefore include a variable corresponding to the difference in parameter measure between the current state (angle/force at TMS) and end-state (maximal parameter measure observed in a trial), i.e. force prediction error or angle prediction error.

We used Bayesian model comparison to investigate whether MEP amplitude modulation is best explained by motor commands or proprioceptive prediction errors. We developed four distinct Bayesian linear mixed models with MEP as the dependent variable. The models were different based on the inclusion of either force/angle at time of TMS, representing optimal control theory, or force/angle prediction error, representing active inference. The prediction error is calculated as the difference between force/angle at TMS and maximal force/angle in the trial. Importantly, for all models we included two electromyography (EMG) measures of muscle contraction since increased EMG facilitates MEP amplitude^7^. Mean EMG 55-5ms prior to time of TMS stimulation (EMG at TMS) and 50-50 ms prior to time of maximal parameter measure were included as variables (mean EMG at max angle/force). We further included subject as a random effect with intercept, since MEP amplitudes vary between subjects, and all continuous variables were standardized per subject. The two force models were compared and the two angle models were compared (Table 1). For an indication of the model fit, we assessed the proportion of MEP amplitude variance explained by the models by estimating bayesian R-squared.

**Table 1.** Bayesian model comparisons. The force optimal control theory model was compared with the force active inference model using Bayesian model comparison, angle n MEPs = 8976, force n MEPs = 3173. See parameter estimates, estimate error and credibility intervals (CI) in Extended data table 1.

|  |  |
| --- | --- |
| Force optimal control theory | Subject + EMG at TMS + EMG at max force + force at TMS |
| Force active inference | Subject + EMG at TMS + EMG at max force + force prediction error |
| Angle optimal control theory | Subject + EMG at TMS + EMG at max angle + angle at TMS |
| Angle active inference | Subject + EMG at TMS + EMG at max angle + angle prediction error |

For both force and angle, we found extreme evidence (Bayes factor > 100) in favor of the optimal control theory model over the active inference model in explaining MEP amplitude modulations when correcting for EMG (force BF mean = 5.69*10^33^, sd = 3.33*10^31^ (sd is 0.59% of the BF mean), angle BF mean = 1.41*10^9^, sd = 1.35*10^7^ (sd is 0.96% of the BF mean)). The proportion of MEP amplitude variance explained by the optimal control theory was R^2^ force estimate = 0.208 and R^2^ angle estimate = 0.0422, indicating that the optimal control theory models do in fact have some predictive power on MEP variability.

Our findings support optimal control theory. MEP amplitude modulations reflect a motor command rather than a proprioceptive prediction error. These conclusions are based on our interpretation of the theories – that the current parameter measure represents a motor command in optimal control theory and that the difference between the current and maximal parameter measures represents the proprioceptive prediction error in active inference. There is conceptual and explanatory flexibility in both theories.

In optimal control theory, the motor command that represents the ‘current state’ could be understood differently than as movement parameter measures. Our interpretation is in line with Shadmehr (2010, pp. 93-94): “For example, if the hand is moving and the brain can predict the current location of the hand and its velocity, then the motor commands should reflect this predicted state”. Here, the ‘predicted state’ is the estimation of the current state created from the inverse model (Fig. 1). The current parameter measures reflected in the MEP amplitude modulation does not necessarily imply direct causality between the motor command and the movement parameters. However, it does suggest a functional linkage between the two.

We built the active inference model based on Adams et al (2013)^4^ and Floegel et al (2023) according to which “The translation from desired perceptions to movements occurs via proprioceptive prediction errors in the spinal cord or brainstem that are quenched by classic reflex arcs”^8^, page 316. If the proprioceptive prediction represents intended perceptual state (max parameter measure) and ascending sensory reafference reflects current proprioceptive state (parameter measure at TMS), then the prediction error propagating through αMN must represent the difference between the two (max parameter measure – parameter measure at TMS). Given this interpretation, the active inference theory is confronted with the explanatory problem that the current parameter level is reflected in the MEP amplitude to a higher degree than is proprioceptive prediction error.

It could be argued that the prediction error does not represent the difference in parameter measures, but rather the error in muscle length, golgi tendon load, etc. But then, why would the M1 be engaged in encoding movement parameters if this signal is not used further down? A rich experimental literature^9–14^, including our own study^15^, have already established this claim.

We do not expect to settle the discussion with this brief communication, nor do we want to dismiss active inference. Our results support optimal control theory over active inference in this experiment. However, there could be other ways to test the theories and indeed other ways to interpret the theories. We encourage proponents of optimal control theory and active inference to specify how their respective theories would be reflected in measurable parameters such as CSE, movement related cortical potentials, motor neuronal firing rate, EMG frequency profile, muscle synergy patterns, state space manifolds, H-reflex, or any other testable quantity. Finding measurable differences might be a challenge. The theories seem very different theoretically but peculiarly similar when sketching the architecture of the testable models.

## Methods

Another manuscript with a different research question and scope is based on the same data^15^. We also obtained data from a velocity task, which was only relevant for the prior study and not presented here.

### Participants

We included 40 healthy participants the study, of which 33 completed (21 female, mean age 29.12, age range 20-45) (3 were excluded because of too high distance between coil and head, 2 did not complete all tasks, 1 terminated the experiment, 1 participant experienced that TMS stimulated their migraine hotspot). We recruited participants through forsøgsperson.dk and University of Copenhagen. We obtained written informed consent for participation and for data collection. We performed TMS in accordance with TMS safety guidelines^17^, and excluded participants with contraindications to TMS safety based on a safety questionnaire developed from Rossi 2009^18^. The study was performed in accordance with the declaration of Helsinki and data was processed and stored according to EU general data protection regulation and guidelines approved by UCPH (2004334 – 4242). The protocol was approved by The Capital Region of Denmark (H-21061035). Participants were compensated with 150 DDK/hour. 2 participants were ambidextrous the rest were right-handed when assessed with the Edinburgh handedness inventory^19^ (83 +/− 22 (mean +/− 1 sd)). Other questionnaires were performed, with relevance to other studies.

### Experimental Design

This study tested the plausibility of two theories, active inference, and predictive coding, by investigating whether the MEP amplitude best represents a motor command or a prediction error in angle and force index finger movement TMS tasks. We also performed a velocity task reported elsewhere^15^, which did not allow a comparison between current and max velocity and is therefore not included in this study.

For both tasks, participants were seated in front of a table with both arms resting horisontally on the table. The right forearm was in a semiprone position, palm facing medially and index finger straight. In the force and angle tasks, the participants aim for a parameter level (force level 1, 3 or 5, angle level 5 or 7). TMS stimulation occurs during the movement at different force/angle levels. This ensures variability in force/angle levels at time of stimulation, and variability in the distance from force/angle levels at TMS to the intended force/angle level. This allows us to investigate which the MEP amplitude reflects to a higher extend: a motor command (force/angle at TMS stimulation) or a prediction error (the difference between current force/angle and intended force/angle). Taken together, for each trial we obtained an MEP amplitude and measures of the parameter (force/angle): 1) the parameter level at time of TMS stimulation, and 2) the difference between the parameter level at time of TMS stimulation and the max parameter level during the trial. Task order was randomized between participants.

### Preparations

Participants performed the experiment in a 6-7 hour visit over one day (two days for one participant). We obtained informed consents and carried out the TMS safety questionnaire and other questionnaires relevant to another study. Electromyography (EMG) electrodes were applied over flexor digitorum profundus (FDP), first dorsal interosseous muscle FDI, and extensor digitorum communis (EDC). Maximal voluntary contraction (MVC) of the right-hand index finger and the maximal compound muscle potential (Mmax) from FDP were measured. Participants had their arm in a semiprone position, palm facing medially and index finger straight at MVC measurement. They pressed three seconds with max force two times, with a 90 second interval between presses. The maximally produced force level in either press was defined as their MVC. We identified TMS stimulation hotspot and motor threshold (MT). During both tasks, visual feedback was prevented by a cloth (force) or box (velocity). Breaks were given when asked for.

### Force task

In the force task, participants pressed the right index finger isometrically against a force transducer in the horizontal direction (Fig. 2). Participants’ fingers were oriented straight ahead, with their position maintained by two supporting sticks, allowing for lean support (Fig. 1). The task required participants to exert force at designated levels—1, 3, or 5—on a 1-10 scale, where 10 represents their maximum voluntary contraction (MVC). These levels correspond to 10%, 30%, and 50% of their MVC, respectively. To minimize fatigue, level 5 was the highest level. Following MVC determination, participants received verbal guidance on achieving force levels 1, 3, and 5, 2-3 times. After this initial guidance, participants had to depend solely on their sensorimotor perception, as no further external feedback was provided.

Importantly, TMS stimulations were applied at force levels relative to participants’ MVC. For force level 1, we expected participants to perform approximately 10% of MVC. Thus, we stimulated half of 10% MVC (5% MVC). For force level 3, we expected participants to perform approximately 30% MVC. We stimulated at either 5% or 15% MVC. For force level 5, we expected participants to perform approximately 50% MVC. During this condition, we stimulated at either 5%, 15%, or 25% MVC. This setup allowed us to obtain MEP measures while we vary force level at TMS stimulation, and the difference between force level at TMS stimulation and max force level in that trial.

During each trial, participants were presented with two distinct auditory tones. The pitch of the initial tone served as an indicator of the target force level: a low, middle, or high pitch corresponded to force levels 1, 3, and 5, respectively. This was followed by a ‘go-tone’, signaling the commencement of the action, which occurred after a variable inter-tone interval. On average, the interval between single TMS stimuli in the force task was 6.0 seconds. Participants completed 5-7 blocks, each lasting 6 minutes (the final block was sham condition), with an average of 90 force trials and 59 sham trials per participant. During sham trials, the TMS coil was positioned such that one wing was perpendicular to the participant’s head.

### Angle task

In the angle task, the right arm and hand were positioned as in the force task, but without the force transducer. Movements of the right index finger were measured using a fiber optic goniometer (Model S720, Measurand Inc., New Brunswick, Canada). The starting position was defined as the index finger pointing directly ahead, designated as angle 0. Participants were trained to consistently return to this initial position. Maximum flexion at the metacarpophalangeal (MCP) joint was identified as angle 9, ensuring no bending occurred at the proximal interphalangeal joint. Within this framework, participants were instructed to move their fingers to either level 5 or 7 in each trial. Prior to the start of the task, levels 5 and 7 were guided 2-3 times using verbal feedback, after which participants received no further external guidance.

Importantly, TMS stimulations were applied at angle levels relative to participants’ angle 9. Angle 0 and 9 were measured, from which a 0-9 scale was calculated. For angle level 5, TMS stimulations were applied at angle 2 or 4. For angle 7, TMS stimulations were applied at angle 2, 4, or 6. This setup allowed us to obtain MEP measures while we vary angle at TMS stimulation, and the difference between angle at TMS stimulation and max angle in that trial.

As in the force task, each trial involved the presentation of two auditory tones. The initial tone, varying in pitch between low and high, signified the target angle level 5 or 7, which was then succeeded by a ‘go-tone’. Participants completed 5-7 blocks, each lasting 6 minutes, with the final block serving as a sham condition. On average, each participant undertook 272 TMS trials and 66 sham trials. The average TMS inter-stimulus-interval during the experiment was 5.3 seconds. Notably, the force task involved an isometric contraction, whereas the angle task required a concentric contraction and hence dynamic movements.

### Data acquisition

The experimental setup was controlled by Spike2 (version 7.10, CED, Cambridge, UK) controlling both the transcranial magnetic stimulation (TMS) and auditory signals via a 1401 Micro Mk II analog-to-digital (AD) converter (Cambridge Electronic Design (CED), Cambridge, UK). Data from the electromyography (EMG), fiber optic goniometer, and force transducer (UU2 load cell, DN-AM310 amplifier, Dacell, Seoul, Korea) were acquired at a sampling rate of 2000 Hz using the CED system. EMG signals were amplified 20 to 100 times (accounted for in the analysis) by a custom-built differential amplifier, developed in the Panum Institute electronics lab, Copenhagen, DK, and were band-pass filtered within a range of 5-1000Hz. All collected data was securely stored on a protected drive.

Surface EMG recordings were obtained from the FDP, FDI, and EDC. This was achieved using Ag/AgCl surface electrodes (22 x 30 mm, Ambu A/S BlueSensor N-10-A/25 ECG electrodes, Ballerup, Denmark) arranged in a belly-tendon montage. The active and reference electrodes were positioned next to each other on the muscle belly, while the ground electrode was secured on the olecranon of the right ulnar bone. To enhance conductivity, electrode gel (Signa Gel, Parker Laboratories, Inc., Fairfield, New Jersey 07004, USA) was applied. Prior to electrode placement, participants’ skin was prepared using fine sandpaper and sanitizer ^20^.

Mmax was elicited through stimulation of the median nerve proximal to the elbow, with recordings from the FDP muscle using a DS7A stimulator (Digitimer Ltd, Welwyn Garden City, England). Although Mmax was recorded, we opted to normalize EMG data relative to the MVC. This approach was chosen because it has been argued that it more accurately reflects voluntary muscle activation^21^, and because Mmax measurements in some participants were not reliable.

### TMS stimulation and MEP measurements

Participants underwent one single-pulse TMS stimulation over the hand-M1 per trial, using a Magstim Rapid2 stimulator (The Magstim Company, Whitland, Dyfed, UK) and a custom-made batwing figure-of-8 shaped coil (biphasic, initial AP direction followed by PA direction, external coil diameter 115 mm). The coil, positioned horizontally over M1, was angled at 45° to the coronal and sagittal planes. Resting motor threshold (rMT) was determined by gradually increasing intensity of stimulator output until TMS induced MEPs (>50μV) in 5 out of 10 trials ^20^. Stimulation intensity in experiments was 120% of rMT with an eight second inter-stimulus interval. The FDP motor hotspot was located lateral to the medial fissure, with the hotspot identified as the point of largest and most consistent MEPs in the FDP. Precision in coil placement (below ∼3 mm from the identified hotspot) was ensured by experimenter, using a BrainSight® neuronavigational system (Rogue Research Inc, Montreal, Canada), and a mechanical holder maintained coil position and thus stimulation of the hotspot. For sham trials, the coil was positioned with one wing perpendicular to the participant’s head.

### Data analysis and statistics

#### EMG analysis

Raw EMG data was corrected for amplification (x20-100), zero-mean centered, smoothed across 80 samples (40ms), normalized to EMG at MVC, and rectified. MEP amplitude, derived from zero-mean centered raw EMG, was normalized against maximum EMG during MVC. MEPs with amplitude below three standard deviations (sd) from background EMG levels were excluded. Analysis was confined to FDP MEPs although we also have FDI and EDC measures. EMG at TMS stimulation is mean smoothed, normalized, and rectified EMG 55-5ms pre-TMS. EMG at max force/angle is mean smoothed, normalized, and rectified EMG 50-0ms pre-max force/angle.

#### Testing which model best explains FDP MEP amplitude using Bayesian model comparison

We performed data analysis in MATLAB (version 9.13 (R2022b), The MathWorks Inc, Natick, MA, USA) and statistical analysis was in R (version 2022.12.0, https://www.r-project.org). Models were compared using Bayes factors, we used the bayes_factor function from the ‘brms’^22^ library (version 2.18.0, https://cran.r-project.org/web/packages/brms/index.html) using STAN^23^ in R. The Bayes factor (BF) is the ratio of the probabilities of the data under two competing models. Samples were collected using Markov Chain Monte Carlo (MCMC) sampling, comprising 4000 iterations, including 2000 warm-up phases, across 7 chains, resulting in a total of 14,000 posterior samples. Chain convergence was evaluated using the Rhat statistic, consistently achieving a value of 1.00. One thousand BFs were computed per test to gain a mean and standard deviation of the BF.

For each parameter force and angle, we computed an active inference model and a predictive coding model. The active inference models included the variable *angle/force at TMS*, and the predictive coding models included the variable *error in angle/force* (Table 1). All models included *subject* as a random effect with intercept, *EMG at TMS* and *EMG at objective force/angle*. MEP amplitude and other continuous variables were standardized per participant by subtracting participant mean and dividing with sd. This standardization facilitated the centering of priors around 0, with a sd of 1. The choice of non-conservative priors is to minimize their influence since there are no previous studies to base the priors on.

#### Testing how much variability the variables in the models explain of the data

To assess the proportion of variance in the MEP amplitude that is explained by the models, we estimated Bayesian R-squared for posterior distributions using the bayes_R2 function from the ‘brms’ package in R (Extended data table 2).

## Additional information

### Data availability

The data supporting the findings in this study are available upon reasonable request from the corresponding author IMB. The data are not publicly available yet since the data are currently pseudo-anonymized. We will fully anonymize the data and make it available with Open Access at Open Science Foundation in case of manuscript acceptance.

### Code availability

The code supporting the findings in this study is available upon reasonable request from the corresponding author IMB. We will make it available with Open Access at Open Science Foundation in case of manuscript acceptance.

### Author contributions

Ida Marie Brandt: Conceptualized and designed the study, data collection, data analysis, discussed results. Writing: original draft.

Thor Grünbaum: Obtained funding, conceptualized the study, provided feedback on experimental setup, discussed results. Writing: Review and editing.

Mark Schram Christensen: Obtained funding, conceptualized, and designed the study, supervised data analysis, discussed results. Writing: Review and editing.

## Acknowledgements

The authors would like to thank Mark Wulf Carstensen and Iva Svecová for helping during data collection, Jesper Lundbye-Jensen for contributing to obtain funding, and Jesper Lundbye-Jensen and Anke Ninija Karabanov and Victor Lange for theoretical contributions to experimental setup and Iva Svecová and Valeria Simonelli for contributing feedback during piloting.

## Supplementary Data

**Extended data figure 1.**
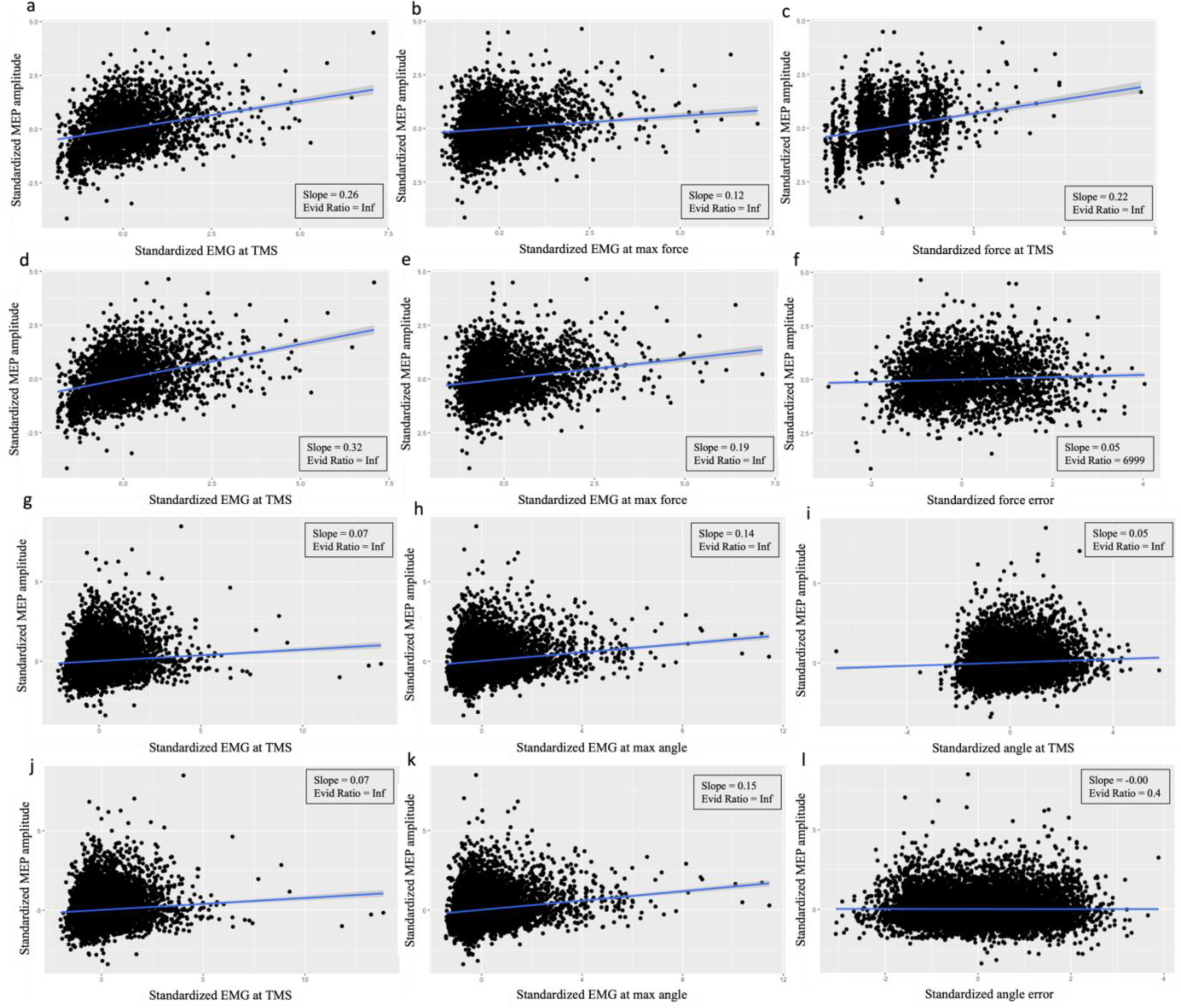
Plot of the conditional effects of the models on standardized MEP amplitude from flexor digitorum profundus. A)-C) Force optimal control theory conditional effects, MEP n = 3173. D)-F) Force active inference conditional effects, MEP n = 3173. G)-I) Angle optimal control theory conditional effects, MEP n = 8976. J)-L) Angle active inference conditional effects, MEP n = 8976. A) Conditional effect of standardized EMG at TMS on standardized MEP from the force optimal control theory model. B) Conditional effect of standardized EMG at max force on standardized MEP from the force optimal control theory model. C) Conditional effect of standardized force level at TMS on standardized MEP from the force optimal control theory model. D) Conditional effect of standardized EMG at TMS on standardized MEP from the force active inference model. E) Conditional effect of standardized EMG at max force on standardized MEP from the force active inference model. F) Conditional effect of standardized force error on standardized MEP from the force active inference model. G) Conditional effect of standardized EMG at TMS on standardized MEP from the angle optimal control theory model. H) Conditional effect of standardized EMG at max angle on standardized MEP from the angle optimal control theory model. I) Conditional effect of standardized angle level at TMS on standardized MEP from the angle optimal control theory model. J) Conditional effect of standardized EMG at TMS on standardized MEP from the angle active inference model. K) Conditional effect of standardized EMG at max angle on standardized MEP from the angle active inference model. L) Conditional effect of standardized angle error on standardized MEP from the angle active inference model.

**Extended data table 1.** Bayesian model estimates of optimal control theory models and active inference models. Variable estimates, estimate error and credibility intervals of the Bayesian linear mixed models from Main table 1.

|  |  | Estimate | Est. Error | CI |
| --- | --- | --- | --- | --- |
| Force motor command | Intercept | 0.00 | 0.02 | [-0.03; 0.03] |
|  | EMG at TMS | 0.26 | 0.02 | [0.23; 0.29] |
|  | EMG at max force | 0.12 | 0.02 | [0.08; 0.15] |
|  | Force at TMS | 0.22 | 0.02 | [0.19; 0.26] |
| Force active inference | Intercept | 0.00 | 0.02 | [-0.03; 0.03] |
|  | EMG at TMS | 0.32 | 0.02 | [0.29; 0.35] |
|  | EMG at max force | 0.19 | 0.02 | [0.16; 0.22] |
|  | Force prediction error | 0.05 | 0.02 | [0.02; 0.08] |
| Angle motor command | Intercept | 0.02 | 0.02 | [-0.02; 0.06] |
|  | EMG at TMS | 0.14 | 0.01 | [0.12; 0.15] |
|  | EMG at max angle | 0.07 | 0.01 | [0.06; 0.09] |
|  | Angle at TMS | 0.05 | 0.01 | [0.04; 0.07] |
| Angle active inference | Intercept | 0.02 | 0.02 | [-0.02; 0.06] |
|  | EMG at TMS | 0.15 | 0.01 | [0.13; 0.16] |
|  | EMG at max angle | 0.07 | 0.01 | [0.06; 0.09] |
|  | Angle prediction error | -0.00 | 0.01 | [-0.02; 0.01] |

**Extended data table 2.** Estimation of MEP variance explained by the models. In the main text we show evidence for optimal control theory over active inference. To assess the proportion of variance in the MEP amplitude that is explained by the models, we estimated Bayesian R-squared for posterior distributions using the bayes_R2 function from the ‘brms’ package in R. The force optimal motor control model estimate = 0.208 and the angle optimal motor control model estimate = 0.0422. We found that the model fit estimates for force models are higher than for the angle models. The Bayesian R2 is a measure of the proportion of variance in the observed data that is explained by the model. Note that the R^2^ estimates cannot be compared between models within the same dataset. An increase R^2^ can therefore not be interpreted as an improved fit^16^.

|  | Estimate | Estimate error | Q2.5 | Q97.5 |
| --- | --- | --- | --- | --- |
| Force simple | 0.167 | 0.0103 | 0.147 | 0.188 |
| Force optimal control theory | 0.208 | 0.0108 | 0.187 | 0.229 |
| Force active inference | 0.171 | 0.0105 | 0.151 | 0.192 |
| Angle simple | 0.0396 | 0.00292 | 0.0340 | 0.0454 |
| Angle optimal control theory | 0.0422 | 0.00299 | 0.0366 | 0.0484 |
| Angle active inference | 0.0396 | 0.00294 | 0.0340 | 0.0454 |

### Bayesian model comparison of active inference models against a model without prediction error

To investigate whether the proprioceptive prediction error does have some explanatory power on MEP amplitude, we tested the active inference force/angle models against a simpler model containing only subject and the two EMG measures (See Extended data table 3). BF active inference force = 8.1 and BF active inference angle = 0.0089. For angle, not only is the model with the prediction error not as good at explaining MEP amplitude as the model with current force/angle measure, it does not add any explanatory power to the MEP amplitude modulations. For force, there was stronger evidence for the active inference model compared to the simple model (BF = 8.1), and the proprioceptive prediction error does thus seem to offer a bit of explanatory power to the MEP amplitude.

**Extended data table 3.**
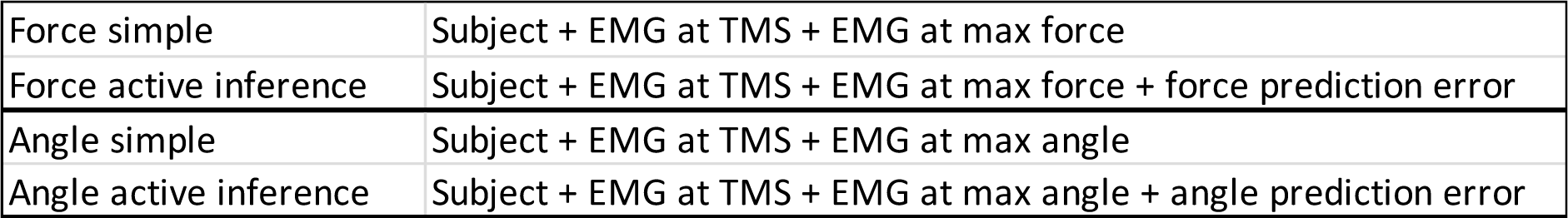
Bayesian model comparisons. The force active inference models and the simple models they are tested against to test whether proprioceptive prediction error offers a degree of explanatory power.

